# A design framework and exemplar metrics for FAIRness

**DOI:** 10.1101/225490

**Authors:** Mark D. Wilkinson, Susanna-Assunta Sansone, Erik Schultes, Peter Doorn, Luiz Olavo Bonino da Silva Santos, Michel Dumontier

## Abstract

“FAIRness” - the degree to which a digital resource is Findable, Accessible, Interoperable, and Reusable - is aspirational, yet the means of reaching it may be defined by increased adherence to measurable indicators. We report on the production of a core set of semi-quantitative metrics having universal applicability for the evaluation of FAIRness, and a rubric within which additional metrics can be generated by the community. This effort is the output from a stakeholder-representative group, founded by a core of FAIR principles’ co-authors and drivers. We now seek input from the community to more broadly discuss their merit.

## Comment

The FAIR Principles[1] provide guidelines for the publication of digital resources such as datasets, code, workflows, and research objects, in a manner that makes them Findable, Accessible, Interoperable, and Reusable (FAIR). The Principles have rapidly been adopted by publishers, funders, and pan-disciplinary infrastructure programmes and societies. The Principles are aspirational, in that they do not strictly define how to achieve a state of “FAIRness”, but rather they describe a continuum of features, attributes, and behaviors that will move a digital resource closer to that goal. This ambiguity has led to a wide range of interpretations of FAIRness, with some resources even claiming to already “be FAIR”! The increasing number of such statements, the emergence of subjective and self-assessments of FAIRness[2, 3], and the need of data and service providers, journals, funding agencies, and regulatory bodies to qualitatively or quantitatively evaluate such claims, led us to self-assemble and establish a FAIR Metrics group (http://fairmetrics.org) to pursue the goal of defining ways to measure FAIRness.

As co-authors of the FAIR Principles and its associated manuscript, founding this small focus group was a natural and timely step for us, and we foresee group membership expanding and broadening according to the needs and enthusiasm of the various stakeholder communities. Nevertheless, in this first phase of group activities we did not work in isolation, but we gathered use cases and requirements from the communities, organizations and projects we are core members of, and where discussions on how to measure FAIRness have also started. Our community network and formal participation encompasses generic and discipline-specific initiatives, including: the Global and Open GO-FAIR (http://go-fair.org), the European Open Science Cloud (EOSC; (https://eoscpilot.eu), working groups of the Research Data Alliance (RDA; https://www.rd-alliance.org) and Force11 (https://www.force11.org), the Data Seal of Approval[4], Nodes of the European ELIXIR infrastructure (https://www.elixir-europe.org), projects under the USA National Institute of Health (NIH)’ Big Data to Knowledge Initiative (BD2K) and its new FAIR Data Com-mons Pilot programmes (https://commonfund.nih.gov/bd2k/commons). In addition, via the FAIRsharing network and advisory board (https://fairsharing.org), we are also connected to open standards-developing communities and data policy leaders, and also editors and publishers, especially those very active around data matters, such as: Springer Nature’s *Scientific Data*, *Nature Genetics* and *BioMedCentral*, *PloS Biology*, *The BMJ*, Oxford University Press’s *GigaScience*, *F1000Research*, *Wellcome Open Research*, Elsevier, EMBO Press and Ubiquity Press.

The converging viewpoints on FAIR metrics and FAIRness, arising from our information-gathering discussions with these variety of communities and stakeholders groups, can be summarized as it follows:

- Metrics should address the multi-dimensionality of the FAIR principles, and encompass all types of digital objects.
- Universal metrics may be complemented by additional resource-specific metrics that reflect the expectations of particular communities.
- The metrics themselves, and any results stemming from their application, must be FAIR.
- Open standards around the metrics should foster a vibrant ecosystem of FAIRness assessment tools.
- Various approaches to FAIR assessment should be enabled (e.g. self-assessment, task forces, crowd-sourcing, automated), however, the ability to scale FAIRness assessments to billions if not trillions of diverse digital objects is critical.
- FAIRness assessments should be kept up to date, and all assessments should be versioned, have a time stamp, and be publicly accessible.
- FAIRness “grade”, presented as a simple visualization, will be a powerful modality to inform users and guide the work of producers of digital resources.
- The assessment process, and the resulting FAIRness grade, should be designed and disseminated in a manner that positively incentivizes the providers of digital resources; i.e., they should view the process as being fair and unbiased, and moreover, should benefit from these assessments and use them as opportunity to identify areas of improvement.
- Governance over the metrics, and the mechanisms for assessing them, will be required to enable their careful evolution and address valid disagreements.

Here we report on the framework we have developed, which encompasses the first iteration of a core set of FAIRness indicators that can be objectively measured by a semi-automated process, and a template that can be followed within individual scholarly domains to derive community-specific metrics evaluating FAIR aspects important to them.

From the outset, the group decided that it would focus on FAIRness for machines - i.e., the degree to which a digital resource is findable, accessible, interoperable, and reusable without human intervention. This was because, by and large, most digital resource are already FAIR for people, and in any case this is not something that can be objectively measured as it will often depend on the experience and prior-knowledge of the individual. We further agreed on the qualities that a FAIR metric should exhibit. A good metric should be:

- Clear: anyone can understand the purpose of the metric
- Realistic: it should not be unduly complicated for a resource to comply with the metric
- Discriminating: the metric should measure something important for FAIRness; distinguish the degree to which that resource meets that objective; and be able to provide instruction as to what would maximize that value
- Measurable: the assessment can be made in an objective, quantitative, machine-interpretable, scalable and reproducible manner, ensuring transparency of what is being measured, and how.
- Universal: The metric should be applicable to all digital resources.

The goal of this working group was to derive at least one metric for each of the FAIR sub-principles, that would be universally applicable to all digital resources in all scholarly domains. We recognized, however, that what is considered FAIR in one community may be quite different from the FAIRness requirements or expectations in another community - different community norms, standards, and practices make this a certainty! As such, our approach took into account that the metrics we derived would eventually be supplemented by individual community members through the creation of domain-specific or community-specific metrics. With this in mind, we developed (and utilized) a template for the creation of metrics (Table 1), that we suggest should be followed by communities who engage in this process.

**Table 1:**
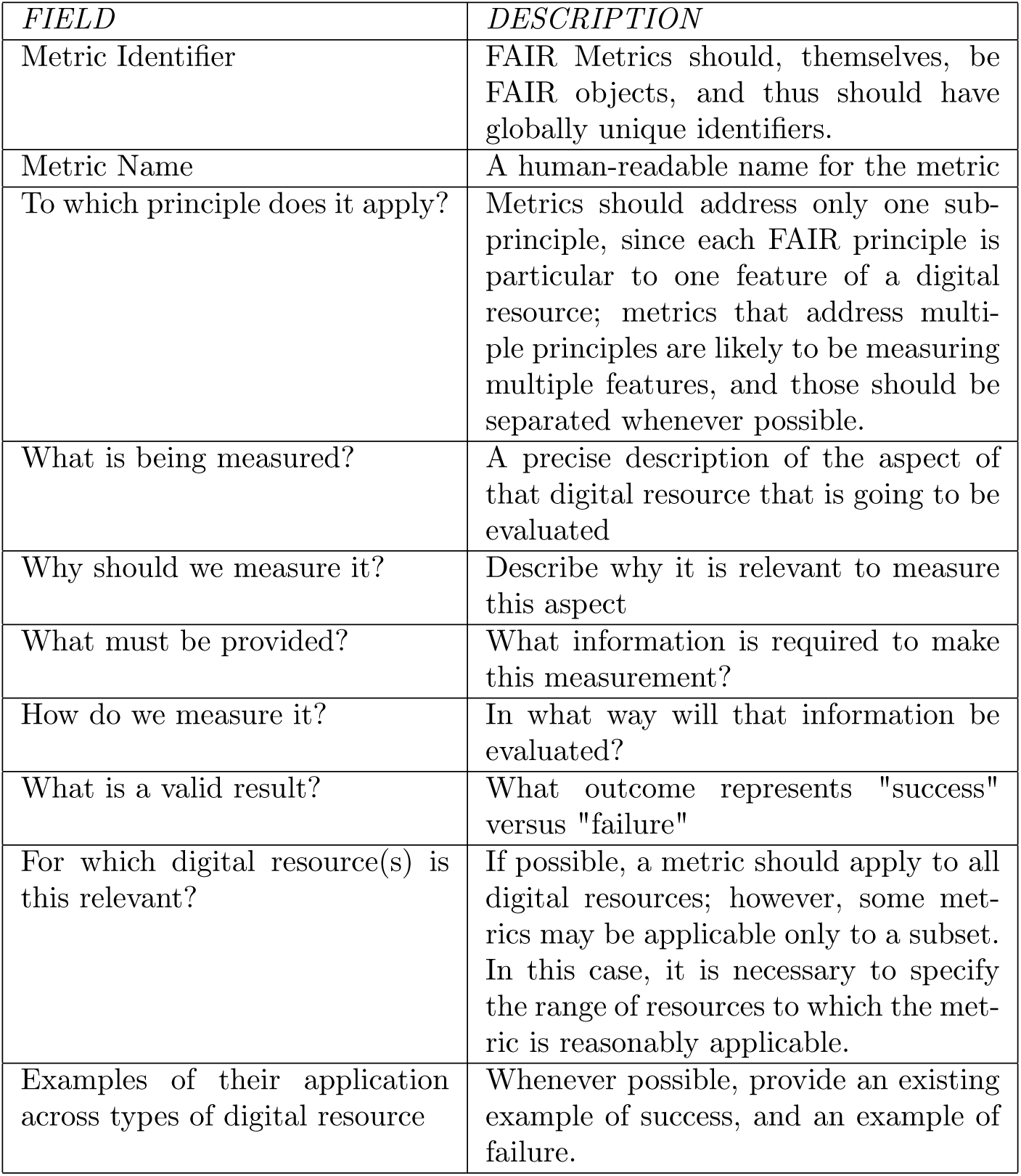
The template for creating FAIR Metrics

| *FIELD* | *DESCRIPTION* |
| --- | --- |
| Metric Identifier | FAIR Metrics should, themselves, be FAIR objects, and thus should have globally unique identifiers. |
| Metric Name | A human-readable name for the metric |
| To which principle does it apply? | Metrics should address only one sub-principle, since each FAIR principle is particular to one feature of a digital resource; metrics that address multiple principles are likely to be measuring multiple features, and those should be separated whenever possible. |
| What is being measured? | A precise description of the aspect of that digital resource that is going to be evaluated |
| Why should we measure it? | Describe why it is relevant to measure this aspect |
| What must be provided? | What information is required to make this measurement? |
| How do we measure it? | In what way will that information be evaluated? |
| What is a valid result? | What outcome represents "success" versus "failure" |
| For which digital resource(s) is this relevant? | If possible, a metric should apply to all digital resources; however, some metrics may be applicable only to a subset. In this case, it is necessary to specify the range of resources to which the metric is reasonably applicable. |
| Examples of their application across types of digital resource | Whenever possible, provide an existing example of success, and an example of failure. |

The outcome of this process was 14 exemplar universal metrics covering each of the FAIR sub-principles. The metrics request a variety of evidence from the community, some of which may require specific new actions. For instance, digital resource providers must provide a publicly accessible document(s) that provides machine-readable metadata (FM-F2, FM-F3) and details their plans with respect to identifier management (FM-F1B), metadata longevity (FM-A2), and any additional authorization procedures (FM-A1.2). They must ensure the public registration of their identifier schemes (FM-F1A), (secure) access protocols (FM-A1.1), knowledge representation languages (FM-I1), licenses (FM-R1.1), provenance specifications (FM-R1.2). Evidence of ability to find the digital resource in search results (FM-F4), linking to other resources (FM-I3), FAIRness of linked resources (FM-I2), and meeting community standards (FM-R1.3) must also be provided. The current metrics are available for public discussion at the FAIR Metrics GitHub, with suggestions and comments being made through the GitHub comment submission system (https://github.com/FAIRMetrics). They are free to use for any purpose under the CC0 license. Versioned releases will be made to Zenodo as the metrics evolve, with the first release already available for download[5]

Finally, we are in the process of prototyping a framework for the automated evaluation of the core metrics, leveraging on a core set of existing work and resources that will progressively become part of an open ecosystem of FAIR-enabled (and enabling) tools. Each metric will be associated with a Web interface capable of evaluating that metric, which is itself described in a FAIR manner using the smartAPI[6] specification. FAIRsharing, will provide source information on metadata, identifier schemas and other standards, which are core element to many metrics. A “FAIR Accessor”[7] will be used to publish groups of metrics together with metadata describing, for example, the community to which this set of metrics should be applied, the author of the metrics set, and so on. An application will discover an appropriate suite of metrics, gather the information required by each metric’s smartAPI (through an automated mechanism or through a questionnaire), and then execute the metric evaluation. The output will be an overall score of FAIRness, a detailed explanation of how the score was derived (inputs/outputs for each metric) and some indication of how the score could be improved. Anyone may run the metrics evaluation tool in order to, for example, guide their own FAIR publication strategies; however, we anticipate that community stakeholder organizations and other agencies may also desire to run the evaluation over critical resources within their communities, and openly publish the results. For example, FAIRsharing will also be one of the repositories that will store, and make publicly available, FAIRness grade assessments for digital resources evaluated by our framework, using the core set of metrics.

Finally, we believe that increasing the FAIRness of digital resources maximizes their reuse, and that the availability of an assessment provides feedback to content creators about the degree to which they enable others to find, access, interoperate-between and reuse their resources. We note, however, that the FAIR-compliance of an resource is distinct from its impact. Digital resources are not all of equal quality or utility, and the size and scope of their audience will vary. Nevertheless, all resources should be maximally discoverable and reusable as per the FAIR principles. While this will aid in comparisons between them, and assessment of their quality or utility, we emphasize that metrics that assess the *popularity* of a digital resource are not measuring its FAIRness. With this in-mind, and with a template mechanism in-place to aid in the design of new metrics, we now open the process of metrics creation for community participation. All interested stakeholders are invited to comment and/or contribute via the FAIR Metrics GitHub site!

## Acknowledgements

We thank all colleagues with whom we discussed the creation and application of the FAIR Metrics; S.-A.S. wants to thank especially Myles Axton, Jennifer Boyd, Helena Cousijn, Scott Edmunds, Emma Ganley, Andrew Hufton, Rebecca Lawrence, Thomas Lemberger, Varsha Khodiyar, Robert Kiley, Michael Markie and Jonathan Tedds for their prospective on the metrics as journal editors and publishers, and their contribution to FAIRsharing. We thank the NBDC/DBCLS BioHackathon series where many of these metrics were designed. The authors received no specific funding for this work, but they want to acknowledge funds supporting them and their research activities. M.D.W. is supported, in part, by the Ministerio de Economía y Competitividad, Spain (TIN2014-55993-RM), and the Dutch Techcenter for Life Sciences. S.-A.S. is funded by grants from the UK BBSRC and Research Councils (BB/L024101/1, BB/L005069/1), EU (H2020-EU.3.1, 634107,H2020-EU.1.4.1.3, 654241, H2020-EU.1.4.1.1, 676559), IMI (116060), and NIH (U54 AI117925, 1U24AI117966-01, 1OT3OD025459-01, 1OT3OD025467-01, 1OT3OD025462-01), which in part also contribute to the FAIRsharing resource. M.D. is funded through several NIH grants (1OT3OD025467-01, 1OT3HL142479- 01, 1OT3TR002027, 5U01HG008473-03) with an emphasis on data sharing.

All authors contributed to the creation of the FAIR Metrics, and the production of this commentary article.

## References

[1] Mark D. Wilkinson, Michel Dumontier, IJsbrand Jan Aalbersberg, Gabrielle Appleton, Myles Axton, Arie Baak, Niklas Blomberg, Jan-Willem Boiten, Luiz Olavo Bonino da Silva Santos, Philip E. Bourne, Jildau Bouwman, Anthony J. Brookes, Tim Clark, Mercè Crosas, Ingrid Dillo, Olivier Dumon, Scott Edmunds, Chris T. Evelo, Richard Finkers, Alejandra Gonzalez-Beltran, Alasdair J.G. Gray, Paul Groth, Carole Goble, Jeffrey S. Grethe, Jaap Heringa, Peter A.C’t Hoen, Rob Hooft, Tobias Kuhn, Ruben Kok, Joost Kok, Scott J. Lusher, Maryann E. Martone, Albert Mons, Abel L. Packer, Bengt Persson, Philippe Rocca-Serra, Marco Roos, Rene van Schaik, Susanna-Assunta Sansone, Erik Schultes, Thierry Sengstag, Ted Slater, George Strawn, Morris A. Swertz, Mark Thompson, Johan van der Lei, Erik van Mulligen, Jan Velterop, Andra Waagmeester, Peter Wittenburg, Katherine Wolstencroft, Jun Zhao, and Barend Mons. The FAIR Guiding Principles for scientific data management and stewardship. Scientific Data, 3:160018, mar 2016.

[2] Dunning, A C and De Smaele, M M E and Böhmer, J K. Evaluation of data repositories based on the FAIR Principles for IDCC 2017 practice paper. (doi:10.4121/uuid:5146dd06-98e4-426c-9ae5-dc8fa65c549f) TU Delft, 2017.

[3] Cox, S. and Yu, J. OzNome 5-star Tool: A Rating System for making data FAIR and Trustable. In Proceedings of the 2018 eResearch Australasia Conference, Oct. 16-20, Brisbane, Australia, 2017.

[4] Dillo, I. and De Leeuw, L. Data Seal of Approval: Certification for sustainable and trusted data repositories. DANS, handle:20.500.11755/300f8a56-e6cf-48a2-8950-17f4f8842df9, 2014.

[5] Mark D Wilkinson, Susanna-Assunta Sansone, Luiz Olavo Bonino, Erik Schultes, Peter Doorn, and Michel Dumontier FAIRMetrics/Metrics: Initial set of proposed FAIR Metrics for community discussion (Version v1.0.1). Zenodo doi:10.5281/zenodo.1065974. November 24, 2017.

[6] Shima Dastgheib, Trish Whetzel, Amrapali Zaveri, Cyrus Afrasiabi, Pedro Assis, Paul Avillach, Kathleen Jagodnik, Gabor Korodi, Marcin Pilarczyk, Jeff De Pons, Stephan Schürer, Raymond Terryn, Ruben Verborgh, Chunlei Wu, and Michel Dumontier. The smartAPI ecosystem for making web APIs FAIR. In Proceedings of the 16th International Semantic Web Conference ISWC 2017, November 2017.

[7] Mark D Wilkinson, Ruben Verborgh, Luiz Olavo Bonino da Silva Santos, Tim Clark, Morris A Swertz, Fleur D.L. Kelpin, Alasdair J. G. Gray, Erik A. Schultes, Erik M. van Mulligen, Paolo Ciccarese, Mark Thompson, Rajaram Kaliyaperumal, Jerven T. Bolleman, and Michel Dumontier. Interoperability and FAIRness through a novel combination of Web technologies. PeerJ Computer Science, 3(e110), 2017.

